# Alarm barks of male vervet monkeys as displays of male quality

**DOI:** 10.1101/2023.06.08.544199

**Authors:** Lukas Schad, Pooja Dongre, Erica van de Waal, Julia Fischer

## Abstract

Research on the vocal behaviour of non-human primates is often motivated by a desire to understand the origins of semantic communication, which led to a partial separation of this research from ecological-evolutionary approaches. To bridge this gap, we returned to the textbook example of semantic communication in animals, the vervet monkey, *Chlorocebus pygerythrus*, alarm call system, and investigated whether male alarm barks fulfil a dual function of alarming and indicating male quality. Barks are loud calls, produced by adult males, in response to large carnivores. However, since barks occasionally occur in agonistic interactions, we investigated whether barks may also indicate male quality. We recorded natural barking events over 23 months, sampling individual male participation from 45 individuals in six free-ranging groups at the Mawana Game Reserve, KwaZulu-Natal, South Africa. We hypothesized that barking frequency is under intra-sexual selection and predicted that barking frequency would increase with male rank and the degree of male-male competition. We found that the highest-ranking males were more likely to produce barks than lower-ranking males and that the number of daily barking events increased during the mating season. We advocate studying primate communication in its evolutionary context to achieve a comprehensive understanding of call “meaning”.

## 1 Introduction

What do nonhuman primate vocalizations mean? This question has been addressed by two independent research streams. The first is fuelled by the desire to understand the origins of speech [1–4] and thus characterized by the search for parallels between human speech and nonhuman primate (hereafter ‘primate’) vocal behaviour. The second research stream is more ecologically and evolutionarily grounded and focusses on signal function and selective pressures [5–8]. The focus on the parallels with human speech may obstruct a deeper understanding of the principles governing primate vocal communication, however. Here, we return to the classic textbook example of semantic communication in monkeys to show how an integrative approach enriches our conception of the meaning of primate calls.

In response to their main predator categories, vervet monkeys produce structurally distinct vocalizations that allow receivers to avoid predation by using different escape responses [9–11]. One of these alarm call types, the ‘threat-alarm-bark’ [9] or ‘leopard alarm’ [10,11], is only produced by adult males, typically in response to mammalian land predators. These ‘barks’ are high amplitude vocalizations that are audible over long distances [9]. In many primate species, adult males produce comparable vocalizations, commonly known as loud calls [8,12–14]. Primate loud calls are assumed to assist in the defence of resources by mediating intra- and inter-group spacing [14] or alerting group members to the presence of specific threats [15,16] while simultaneously deterring predators in the area [17,18].

Primate loud calls have also received attention as potential targets of sexual selection that may fulfil a function in mate attraction or male-male competition [19–24]. Physiological constraints affecting signal structure or direct energetic costs related to signal production can impose limitations on individual vocal structure and call rate, rendering loud calls potential honest signals advertising male condition and competitive ability [5]. Loud call structure and call usage vary with signaller rank in several primate species, suggesting a link between individual vocal behaviour and competitive ability [23,25–27]. Due to their high amplitudes, barks are perceived by predators, potential mates and male competitors, whose responses may vary with the signal’s implications for their fitness [18,28]. Costs related to call production might result in a ‘quality handicap’, whereby barks could become honest signals of individual competitive ability [5,29]. Such a mechanism has been suggested for the contest barks produced by chacma baboons, *Papio ursinus*, and geladas, *Theropithecus gelada*, during male-male conflicts [23,24,30]. Since loud calls are audible to many different receivers simultaneously, natural and sexual selection may result in more than one function in such signals [21,22,31–33].

Previous studies of male vervet monkey loud calls have focused on the alarm and predator deterrence function of barks and examined variation in call structure and receiver responses to barks in playback experiments. The degree of inter-individual variation in bark usage across contexts and with regard to signaler demographic and social factors has received comparatively little attention [34,35]. Characterizing signal function and identifying the evolutionary mechanisms that allow vocal signals to exert their effects on receivers requires examining variation in signal usage with regard to all aspects of a species’ social ecology.

Vervet monkeys live in multi-male, multi-female groups with female philopatry. Males typically depart from their natal groups around the time of sexual maturity and subsequently keep migrating between groups throughout their lives, typically during the mating season [36,37]. Among adult males, repeated agonistic interactions are common and increase in frequency and severity during the mating season [36,38]. Male relationships are characterized by a dominance hierarchy, yet there is only moderate reproductive skew [38]. Previous reports suggest that barks are occasionally produced during within and between-group aggressive interactions [9,39,40], opening the possibility that they may also function as intra-sexual displays [39].

In this study, we tested two non-mutually exclusive hypotheses: (I) Barks are honest signals of individual competitive ability and under intra-sexual selection. (II) Males adjust their bark production according to the number of potential offspring in the group, which should correlate with their tenure on average [38]. In the first part of this study, we examined how frequently individual males produced barks in group-level events in relation to individual quality, tenure and the degree of male-male competition. We used rank as a proxy for individual quality, as this construct is presumed to reflect male competitive ability. We gauged male-male within-group competition by considering the number of adult males in the group and the adult sex ratio, measured as the number of adult females per male. Lastly, we assumed increased male-male competition during the mating season (yes/no) [36], which is approximately between April and July, based on previous work in this species [36–38] and the distribution of births in our study period, assuming a 165 day gestation period [37,38]. We calculated male tenure as the number of mating seasons an adult male was present in a group, since even partial presence during the mating season can result in reproductive success [38], which could not be measured directly.

If barks were honest signals of individual competitive ability, the likelihood of participating in a barking event should increase with individual quality and the degree of male-male competition. Therefore, we predicted a three-way interaction among rank, mating season and the number of adult males in the group as well as a three-way interaction among rank, mating season, and adult sex ratio. Male calling probability was expected to increase with male rank, with the steepness of the increase being greater with more male competitors, lower ratios of females to males, and when events took place during the mating season. However, if barks were primarily adaptive due to their alarm function, we would expect that males gain differential fitness benefits from alarm calling depending on the number of potential offspring they had in the group. Males with more offspring would be expected to profit more from alarm calling and should thus show higher barking activity. Due to moderate reproductive skew among male vervets [38], we assumed male offspring number to correlate positively with tenure and predicted that individual likelihood of participating in barking events should increase with individual tenure.

In the second part of the study, we investigated whether the daily number of barking events was related to group composition and the mating season. We predicted that, if barks were honest signals of male competitive ability and males used these calls as assessment signals, the daily number of barking events would rise with increased male-male competition. We therefore expected an interaction between season and the number of adult males in the group as well as an interaction between season and adult sex ratio. The daily number of alarm events was predicted to increase during the mating season, with the steepness of the increase being greater for a larger number of males and a lower number of females per male.

## 2 Methods

### 2.1 Study site, study subjects and data collection

Over the course of 23 months (May 2020 – April 2022), 43 trained observers collected data on six habituated groups of free-ranging Vervet monkeys at the Inkawu Vervet Project (28°00.327S, 031°12.348E) in the Mawana Game Reserve, KwaZulu-Natal, South Africa (2213 observation days). All researchers were trained to identify subjects via individually specific morphological features and to collect data with a minimum inter-observer reliability agreement of 80% Cohen’s Kappa [41]. Group composition varied due to birth, mortality, maturation, dispersal and immigration. We collected ad-libitum data on dyadic agonistic interactions between subjects, individual daily presence in group logbooks and participation of individual adult males in barking events. During barking events, we recorded the identity of every adult male producing barks, the number of unidentified male callers, the context of the calling bout, and the duration of the event. An event was determined to be over when calling at the group-level ceased for more than five minutes. Since barks are audible over several hundred meters, we consider it very unlikely that we missed any events while observers were with a group. Maps showing the GPS coordinates of events and estimated audible distances of barks are available in the electronic supplementary material (ESM) in addition to distributions of event contexts, event durations and number of callers in the events (fig. S1 – S5).

We used data on decided dyadic conflicts from all subjects to determine dominance hierarchies based on Elo-ratings [42] and added new individuals joining a group as soon as observers could identify them. We estimated male hierarchies by ranking individual adult males according to their Elo-scores and calculated the proportion of other adult males an individual dominated on any observation day in their respective group. We chose this ranking system to obtain proportional male ranks that were comparable between groups with different numbers of adult males.

### 2.2 Statistical analysis – General procedure

All statistical analysis was conducted in R (v. 4.2.1) [43] using the lme4 package (v. 1.1-30) [44] and the glmmTMB package (v. 1.1.4) [45]. We fitted a binomial and a zero-inflated Poisson model and included all theoretically identifiable random slopes into the models to reduce type I error rate [46] and excluded correlations among random intercepts and slopes when unidentifiable (absolute correlation parameters ≥ .9) [47]. Visual inspection of histograms for every random intercept and slope component supported the assumption that the best linear unbiased predictors for random effects were normally distributed [48]. To ease comparability of model estimates, we z-transformed covariates to a mean of zero and a standard deviation of 1 and dummy coded and centred the factor mating season (Yes/No) in the random effects parts of the models, using ‘outside mating season’ as reference category [49]. We further used the optimizer ‘bobyqa’ in the binomial model to assist model conversion. To rule out collinearity among predictors, we checked variance inflation factors using the R package ‘car’ (v. 3.1.0) [50] on simplified general linear mixed models excluding the random effect structure and the interactions.

For the zero-inflated Poisson model, we checked for overdispersion to avoid potential type I errors and corrected reported standard errors [51]. To prevent ‘cryptic multiple testing’ [52], we compared the full models to null models that only included control predictors and offset terms but the same random effects structure as the full models by using a likelihood ratio test [53]. Model stability was assessed by comparing the estimates of the full model with those obtained from identical models fitted to subsets of the data that excluded the individual levels of the random effects one at a time (function provided by R. Mundry) [54]. We determined 95% confidence intervals with the ‘bootMER’ function of the ‘lme4’ package using 1000 parametric bootstraps. P-values for individual fixed effects and interactions were derived from likelihood ratio tests using the R function ‘drop1’ with a Chi-square test argument [46]. If interactions were not significant (P > 0.05) according to likelihood ratio tests, we fitted reduced models by removing the respective interactions as main effects without changing the random effect structure of the models. Detailed information about model structure, random slopes, model stability and the results of likelihood ratio tests for interactions are available in the supplementary material (table S1 – S3).

### 2.3 Binomial model – Individual calling probability in barking events

We used a Generalized Linear Mixed Model [48] with binomial error structure and logit link function to investigate which predictors influenced male probability to produce barks in a barking event. As fixed effects, we incorporated the covariates male rank (range: 0 – 1), number of adult males in the group (mean ± s.d. = 7.5 ± 3.3; range: 2 to 13), adult sex ratio (number of females per male; mean ± s.d. = 2.0 ± 0.7; range: 1 to 5.5), male tenure (mean ± s.d. = 2.2 ± 1.4 mating seasons; range: 0 to 7) and the factor mating season (yes/no). We included the 3-way interaction among male rank, mating season, and number of adult males, as well as the 3-way interaction among male rank, mating season, and adult sex ratio, while incorporating all lower order terms these encompassed. As random factors, we included individual ID, group ID, event ID and the combination of date and group to control for potential non-independence of different events observed in the same group on the same day. We excluded all events in which unknown males produced barks from the analysis since the barking activity of unknown callers would introduce uncertainty into the binomial response ‘male produced barks (Yes/No)’. We also excluded events from groups that temporarily just had a single resident adult male since the predictor proportional male rank is not defined in single-male groups.

### 2.4 Poisson model – Number of barking events per observation day

To assess whether group composition and the mating season affected the number of barking events recorded per day, we used a Generalized Linear Mixed Model with a Poisson error structure and a log-link function accounting for zero-inflation [45,55]. We included the covariates number of adult males (mean ± s.d. = 5.6 ± 3.1; range: 1 to 13), adult sex ratio (mean ± s.d. = 2.4 ± 1.1; range: 0.7 to 12) and the factor mating season (yes/no) as fixed effects. Further, we added the interaction between mating season and number of adult males as well as the interaction between mating season and adult sex ratio. As a control predictor, we included the covariate group size (mean ± s.d. = 36.6 ± 15.8; range: 5 to 77). To account for observation effort, we added the log-transformed observation time of every observation day as an offset term (mean ± s.d. = 6.9 ± 2.1; range: 2.5 to 14). As a random factor, we included group ID. An initial Poisson model revealed that the observed number of zeros was higher than expected given the model. To account for zero-inflation we incorporated a random intercept in the zero-inflation part of the model, but none of the predictors from the count part of the model. We excluded all days with missing observation times from the analysis.

## 3 Results

### 3.1 Binomial model – Individual calling probability in barking events

In the 23 months, we recorded 886 barking events. After excluding all events with unknown male callers and all events in single male groups, the sample for the model comprised 369 events, yielding 2055 individual data points (with 524 barking participations) from 45 adult males in six groups over 291 group observation days. Collinearity among predictor variables did not pose a problem (all variance inflation factors < 1.7). The fixed effects of the model had an overall impact on the probability to produce barks in an event, as shown by comparing the full and null model (*χ2* = 25.326, *df* = 12, *P* = 0.013). Since likelihood ratio tests indicated that the three-way and two-way interactions were not significant (ESM, table S1 – S2), we removed them and fitted a reduced model. The reduced model revealed no obvious effects of adult sex ratio, mating season and male tenure (table 1). Higher-ranking individuals had a higher probability to engage in barking activity (fig. 1 and table 1). The positive effect of rank on calling probability appeared moderate overall, except for the highest-ranked males in a group, who showed very high calling probability compared to all other ranks (fig. 1). Visual inspection of the model (fig. 1) indicated a poor model fit, suggesting that the behaviour of the highest ranked males in the group predominantly drove the positive effect of rank. We therefore refitted the model excluding the highest ranked males (rank = 1; 23 of 45 males temporarily held the highest rank). While the estimate for rank was still positive, visual inspection of the refitted model corroborated the assumption that the effect of rank was mainly driven by barking activity from the highest ranked males in the group (ESM, fig. S8).

**Figure 1:**
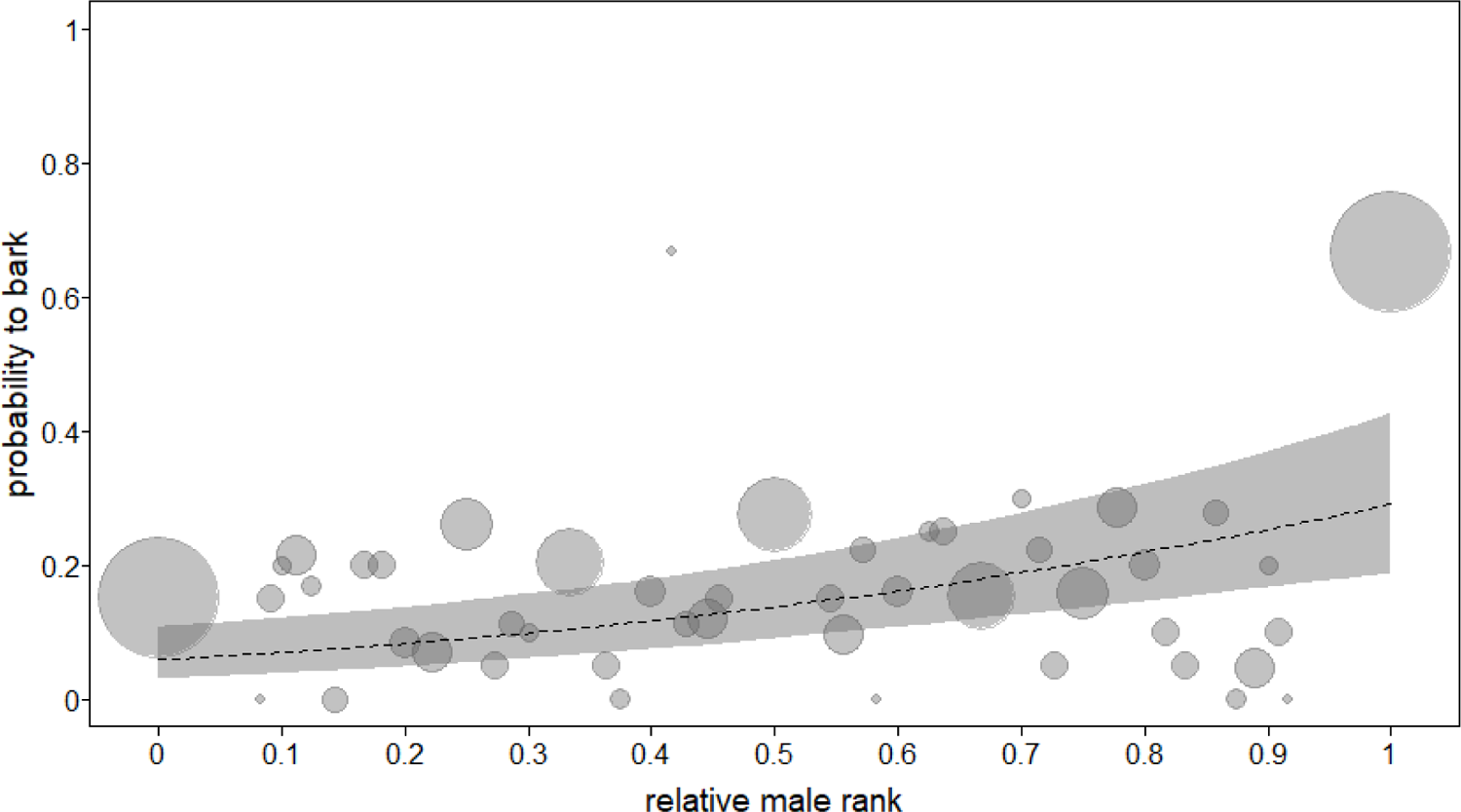
Probability to bark in relation to individual rank (proportion of other males dominated in the group). The plot shows the average barking probability for every rank value, with the circleś area proportional to the frequency that a given rank value occurred (range: 3 to 372). The dashed line depicts the fitted model and the grey shaded area the bootstrapped 95% confidence intervals, with all other predictors being at their mean for covariates and the factor mating season being dummy coded and centred. Higher-ranked individuals were more likely to participate in a barking event (GLMM: n = 2055, *p* = 0.002).

**Table 1:** Model results showing the impact of the fixed effects on the probability to produce barks in an event. Shown are model estimates, standard errors, confidence intervals (CI) and the test results obtained from likelihood ratio tests.

| Term | Estimate | SE | Lower CI | Upper CI | $\chi^2$ | df | P |
| --- | --- | --- | --- | --- | --- | --- | --- |
| **Intercept** | -1.767 | 0.266 | -2.263 | -1.165 |  |  | * |
| **Rank** | 0.667 | 0.144 | 0.363 | 0.941 | 9.589 | 1 | 0.002 |
| **Males** | -0.919 | 0.238 | -1.399 | -0.421 | 10.479 | 1 | 0.001 |
| **MatingSeasonY** | -0.133 | 0.238 | -0.617 | 0.341 | 0.325 | 1 | 0.568 |
| **SexRatio** | 0.022 | 0.139 | -0.272 | 0.315 | 0.024 | 1 | 0.876 |
| **Tenure** | 0.268 | 0.184 | -0.110 | 0.685 | 2.021 | 1 | 0.155 |

\* Not shown due to limited interpretability.

In addition to a positive effect of rank on calling probability, the model revealed that a higher number of adult males in the group was associated with a decreased probability to produce barks in an event (fig. 2A and table 1). The distribution of the number of calling males showed that in most events, only a single male was calling (fig. 2B). Notably, the highest-ranked males were responsible for more than half of the single male calling events and barked in a large proportion of all other events as well (fig. 2B).

**Figure 2:**
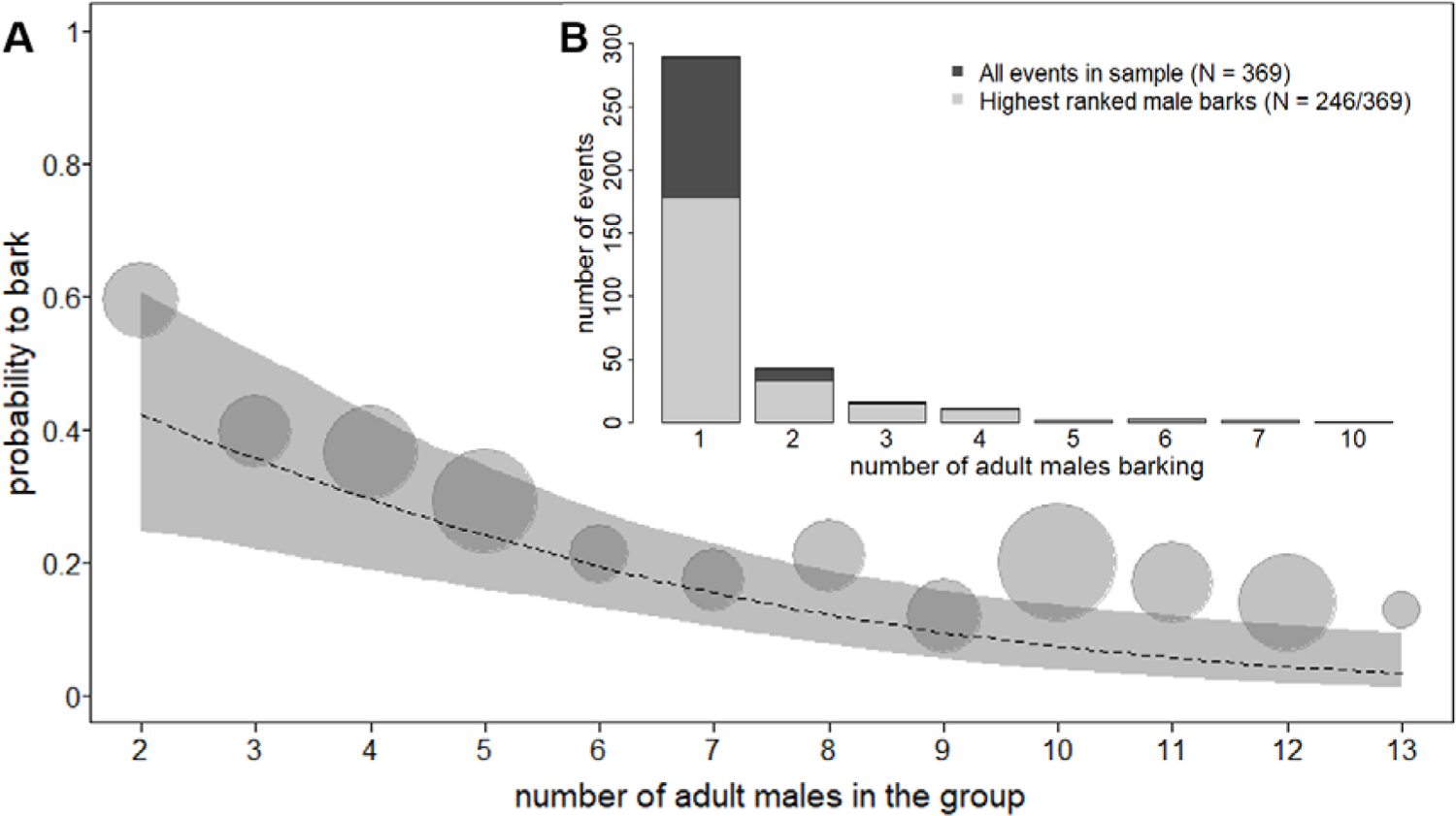
**A** Probability to bark for any given male in relation to the number of males in the group. The plot shows average barking probability, with the circle area proportional to the frequency at which a given number of males occurred (range: 39 to 360). The dashed line depicts the fitted model and the grey shaded area the bootstrapped 95% confidence intervals, with all other predictors at their mean or dummy coded and centred, respectively. Higher numbers of males decrease barking probability (GLMM: n = 2055, *p* = 0.001). **B** Distribution of the total number of callers across analysed barking events (N = 369). The light grey area shows the proportion of events where the highest-ranked male produced barks in the event (N = 246). In most events, only a single adult male called. The highest-ranked males were active in the majority of events.

### 3.2 Poisson model – Number of barking events per observation day

After excluding days with missing observation times, the sample for analysis comprised 1915 group observation days with 821 barking events from six groups. Collinearity among predictor variables could be excluded for ‘mating season’ and ‘sex ratio’ with variance inflation factors < 1.9. We could not exclude potential collinearity for the predictor ‘number of adult males’ and the control predictor ‘group size’ with variance inflation factors < 6.7 and < 5.1, respectively. Since larger groups typically contain more adult males, a positive relationship between the number of adult males in a group and group size can be expected. However, due to fluctuating group composition during the study time, we decided that controlling for group size was required and did not change the model structure. Overdispersion could be ruled out (dispersion parameter = 1.08). The fixed effects had a general impact on the number of group-level barking events per day, as shown by comparing the full and the null model (*χ2* = 20.387, *df* = 5, *P* = 0.001). Subsequent likelihood ratio tests indicated that none of the interactions were significant (ESM, table S3). The reduced model revealed that during the mating season, the number of barking events per day increased (fig. 3 and table 2; also see ESM fig. S6 – S7 for the monthly distribution of barking events and conceptions, respectively). Group size, adult sex ratio and the number of adult males did not significantly affect the number of bark events per day (table 2).

**Figure 3:**
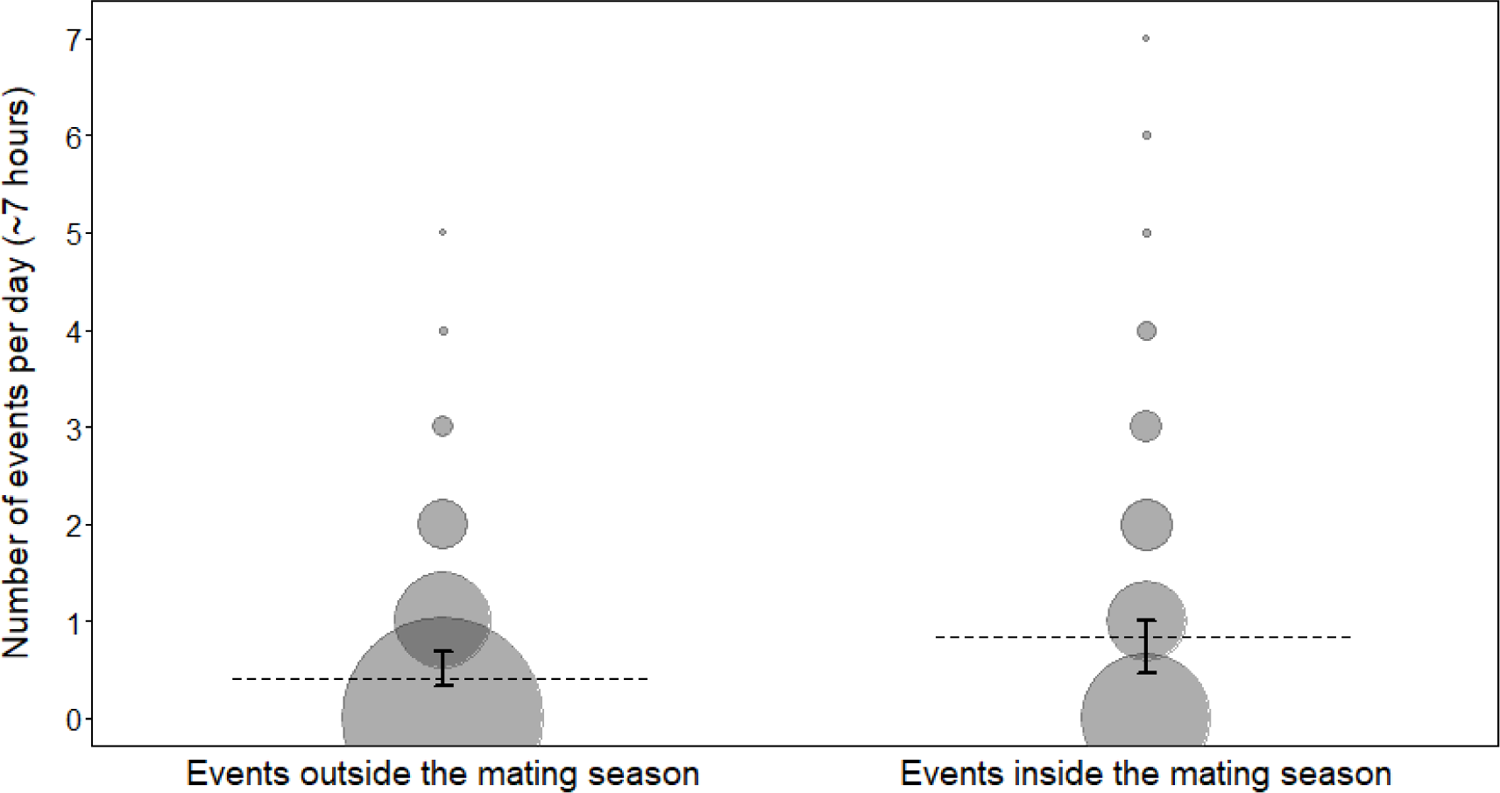
The effect of the mating season on the number of barking events recorded per observation day, for an average observation time of approximately seven hours. The plot shows the daily number of barking events, with the circle area proportional to the frequency of the respective number of events recorded per day (range: 1 to 1358). The dashed grey line depicts the fitted model, and black error bars indicate the bootstrapped 95% confidence intervals, with all other predictors being at their average. Barking events occurred more frequently during the mating season (GLMM: n = 1915, *p* < 0.001).

**Table 2:** Model results show the impact of the fixed effects on the number of barking events per day. Shown are model estimates, standard errors, 95% confidence intervals and the test results obtained from likelihood ratio tests.

| Term | Estimate | SE | Lower CI | Upper CI | $\chi^2$ | $df$ | $P$ |
| --- | --- | --- | --- | --- | --- | --- | --- |
| **Intercept** | -2.658 | 0.095 | -2.876 | -2.490 |  |  | * |
| **MatingSeasonY** | 0.742 | 0.098 | 0.532 | 0.930 | 14.798 | 1 | <0.001 |
| **SexRatio** | 0.042 | 0.084 | -0.139 | 0.186 | 0.274 | 1 | 0.601 |
| **Males** | 0.187 | 0.117 | -0.068 | 0.402 | 2.353 | 1 | 0.125 |
| **GroupSize** | -0.109 | 0.105 | -0.325 | 0.108 | 1.087 | 1 | 0.297 |
| **Zero-inflation-Intercept** | -0.555 | 0.145 | -0.886 | -0.303 |  |  | * |

\* Not shown due to limited interpretability.

## 4 Discussion

The barking probability of adult male vervet monkeys varied with the rank that males held at the time of the event. The highest-ranked males in a group showed comparatively high calling probability, which was likely driving the positive effect of rank. In line with previous studies on chacma baboons and crested macaques, *Macaca nigra* [23,24,27], our results thus support the idea that loud calling behaviour depends on individual rank and indicates caller quality. Contrary to our predictions, an increasing number of males per group decreased individual barking probability, due to the high number of events with only a single caller. Although male vervet barking can thus be viewed as a signal of quality, these calls are not used as displays before fighting to resolve conflicts at lower costs, as is the case in chacma baboons, for instance. We also found no evidence for a relationship between *individual barking probability* and the adult sex ratio, the mating season and the number of potential offspring in the group, as reflected by male tenure [38].

The *number of barking events* recorded in a group per day increased during the mating season, while the adult sex ratio and the number of males per group had no obvious effect. The distribution of births during the study time suggested that most females conceived between May and June, which were the two months with the highest number of barking events in our dataset. As female vervet monkeys are assumed to conceal the time of ovulation from males [56], the increased number of daily barking events during the mating season suggests that males likely responded to the occurrence of mating behaviour with increased barking activity. Increased loud call production in vervet monkeys during the mating season is in line with reports from other species of catarrhine primates, in which males increased rates of loud call production around the time of ovulation [31,32,57].

The positive relationship between high rank and increased barking frequency suggests that individual barking activity reflects male competitive ability or motivation to compete [5,29,58], contradicting the prevalent assumption that barks are predominantly alarm calls that are mostly produced in response to large carnivores. If bark production was costly, either by direct energy expenditure or opportunity costs such as missed foraging time [59], high ranking males could be assumed to incur higher potential costs than other males in the group. Barks may thus be ‘quality handicaps’ [29] and an honest signal of male competitive ability if lower-ranked individuals could not afford the potential costs associated with frequent barking activity. There is evidence in several primate species that male loud call usage varies with individual rank, condition or exhaustion after male-male conflicts, supporting a quality handicap function [24,27,30,57,60]. In longer barking events with more than one caller that may even be triggered by an actual predator, prolonged male barking could thus not only deter predators [17,18], but simultaneously allow males to display stamina to one another, which would speak for a quality handicap. This hypothesis would be supported if rank would also predict individual call rate during events. Currently, only one study has reported that higher ranked male vervet monkeys bark for longer durations compared to lower ranked individuals [34], but the authors were limited to a single experimental leopard model presentation in two groups. However, since male vervet monkeys do not typically engage in ritualistic chases involving signalling contests, and events with a high number of callers are quite rare, the support for a quality handicap mechanism is presently limited.

Instead, the high frequency of events with just one barking male suggests the alternative mechanism of a conventional signal that primarily indicates male motivation to compete [29,61]. Conventional signals are assumed to remain honest even when the costs of signal production are low due to a receiver retaliation rule, which effectively punishes individuals who produce such signals but are not in good enough condition to escalate conflicts into physical fights if receivers attack them [29,62]. The possibility of receiver retaliation would constrain bark production to males in good enough condition to risk costly fights. Importantly, if dominant males would punish lower-ranked males who produce barks, then barking outside of predator events would ensue the risk of provoking a physical fight for all males except the presently highest ranked individual. Following this scenario, we may expect a suppression of frequent barking activity in most males, except for the highest-ranking individuals, which would fit the observed pattern and explain why most barking events only had a single caller. While frequent barking events with just one caller could also be explained by high false alarm rates, it is questionable why false alarms should be more frequent during the mating season and why high-ranking males would have particularly high false alarm rates.

We contend that the mechanisms for a quality handicap signal of male competitive ability or for a conventional signal of motivation to compete, are complementary. Follow-up studies should also aim to clarify whether the acoustic structure of barks is related to caller dominance, body mass or age, as has been found in several primate species [23,27,30,63–65]. Such a relationship would imply that barks might even have index-like properties reflecting caller condition [29].

Whether loud calls initially evolved as alarm calls due to specific predation pressure and subsequently served as substrate for sexual selection [21], or represent an ancestral trait in primates [19] remains unclear. A recent proposal suggested that loud alarm calls of male primates might also constitute inter-sexually selected costly signals, if females were selecting males for providing anti-predation services [22]. In the case of the vervet monkey, we see more support for the assumption that bark production is an intra-sexually selected trait. Firstly, we did not find bark production to be reserved to predator contexts. Secondly, if females would prefer males who bark frequently, we would expect most males to participate in the majority of barking events, which was not the case. Instead, the highest-ranked males frequently barked alone, suggesting that barks are displays indicating caller dominance and barking activity of lower ranked males is suppressed.

The present study adds an important and previously overlooked facet to our understanding of the vervet monkey alarm call system. Despite several reports that calls similar to the well-known alarm calls given to leopards, snakes and eagles may occur in other contexts [9,39,40], the general focus of previous work remained on the acoustic differences between these three alarm call types. By showing that male barks not only occur in predator contexts and that their usage depends on individual rank, we add to the debate about context-specificity in primate calls and the concept of ‘functional reference’ [66]. In addition, our findings raise questions regarding how receivers decide whether they are confronted with alarm or rather display calls. Most likely, receivers are able to factor in individual identity (including the subject’s dominance rank), contextual cues, and season to predict whether a call bout was given in response to a predator or not. The relatively moderate reactions of free-ranging vervets to naturally occurring barking events [59] and variable responses in playback experiments [39,67,68] also support the assumption that receivers may not invariably infer predator presence from the occurrence of barks [69]. Our study thus highlights that the prevailing focus on the potential semantic content of primate calls may divert attention away from other aspects, such as the selective pressures that shape a communication system [31,32].

We suggest that future studies should address long-term ontogenetic changes in bark usage with age, rank, aggression rate, tenure, health status and an individual’s social position within a group. While we presently do not see strong evidence for a specific mechanism that would make barks reliable signals of male competitive ability, future analysis of call rate and call structure may substantiate the interpretation that barks are quality handicaps and under intra-sexual selection. If barks were used as conventional aggressive signals, we would expect that an increase in individual aggression rate should be accompanied by an increase in individual barking frequency as well. We were under the impression that such a pattern was typical for males that had recently acquired the highest rank in their group, but could not formerly test the hypothesis due to a lack of focal data. Comparing bark usage among captive, urban living, and free-ranging populations inhabiting different environments may further reveal whether ecological factors like population-specific predation pressure also affect vocal behaviour in this species.

## Supporting information

Electronic supplementary material

## Ethics

This research adhered to the Association for the Study of Animal Behaviour Guidelines for the Use of Animals in Research [70], the regulations set by the local authority, Ezemvelo KZN Wildlife, as well as the Animal Care Committee at the German Primate Center. We further obtained permission from the owners of the Mawana Game Reserve, the van der Walt family.

## Data accessibility

All data and code are available on OSF as ‘Alarm barks of male vervet monkeys as displays of male quality’ via the following link: https://doi.org/10.17605/OSF.IO/2W7ZH

## Authors’ contributions

LS and JF conceived the study and acquired the funding; LS collected the data, conducted the statistical analyses, prepared the figures and drafted the first version of the manuscript. PD provided the male hierarchy data and assisted with data collection. EW provided the long-term data set, logistics and funding for the field site. All authors edited the manuscript.

## Conflict of interest declaration

We declare to have no competing interests.

## Funding

We gratefully acknowledge funding by the Deutsche Forschungsgemeinschaft (DFG FI707/25-1 – Project Number 428036558) and the Leibniz Association through the Leibniz ScienceCampus Primate Cognition (Seed fund: LSC-SF2018-09). We thank the Swiss National Science Foundation who funded the Inkawu Vervet Project as well as PD and EW salaries (grants to EW: PP00P3_198913 and PP03P3_170624).

## Acknowledgements

We thank the van der Walt family for permission to work at the Mawana Game Reserve. We further thank Michael Henshall and all members of the IVP field team for their help with data collection and invaluable commitment to keeping the IVP field site alive throughout the COVID-19 pandemic. We are particularly grateful to Roger Mundry for statistical teaching and advice and to Robert Seyfarth and Peter Henzi for helpful comments on the manuscript.

## Notes

### Competing Interest Statement

The authors have declared no competing interest.

## References

1. Fitch WT. 2010 The Evolution of Language. Cambridge: Cambridge University Press. (doi:10.1017/CBO9780511817779)

2. Hauser MD, Chomsky N, Fitch WT. 2002 The Faculty of Language: What Is It, Who Has It, and How Did It Evolve? Science (80-.). 298, 1569–1579. (doi:10.1126/science.298.5598.1569)

3. Fischer J. 2017 Primate vocal production and the riddle of language evolution. Psychon. Bull. Rev. 24, 72–78. (doi:10.3758/s13423-016-1076-8)

4. Fedurek P, Slocombe KE. 2011 Primate vocal communication: A useful tool for understanding human speech and language evolution? Hum. Biol. 83, 153–173. (doi:10.3378/027.083.0202)

5. Maynard Smith J, Harper D. 2003 Animal Signals. Oxford: Oxford University Press.

6. Rendall D, Owren MJ. 2013 Communication without meaning or information: abandoning language-based and informational constructs in animal communication theory. In Animal Communication Theory (ed U Stegmann), pp. 151–188. Cambridge: Cambridge University Press.

7. Puts DA et al. 2016 Sexual selection on male vocal fundamental frequency in humans and other anthropoids. Proc. R. Soc. B Biol. Sci. 283, 20152830. (doi:10.1098/rspb.2015.2830)

8. Waser PM, Brown CH. 1986 Habitat acoustics and primate communication. Am. J. Primatol. 10, 135–154. (doi:10.1002/ajp.1350100205)

9. Struhsaker TT. 1967 Auditory communication among vervet monkeys (*Cercopithecus aethiops*). In Social communication among primates. (ed SA Altmann), pp. 281–324. Chicago: University of Chicago Press.

10. Seyfarth RM, Cheney DL, Marler P. 1980 Monkey responses to three different alarm calls: evidence of predator classification and semantic communication. Science (80-.). 210, 801–803.

11. Seyfarth RM, Cheney DL, Marler P. 1980 Vervet monkey alarm calls: Semantic communication in a free-ranging primate. Anim. Behav. 28, 1070–1094.

12. Gautier JP, Gautier A. 1977 Communication in old world monkeys. In How animals communicate (ed TA Sebeok), pp. 890–964. Bloomington: Indiana University Press.

13. Mitani JC, Stuht J. 1998 The evolution of nonhuman primate loud calls: Acoustic adaptation for long-distance transmission. Primates 39, 171–182. (doi:10.1007/BF02557729)

14. Byrne RW. 1982 Primate Vocalisations: Structural and Functional Approaches to Understanding. Behaviour 80, 241–258. (doi:10.1163/156853982X00373)

15. Zuberbühler K, Noë R, Seyfarth RM. 1997 Diana monkey long-distance calls: Messages for conspecifics and predators. Anim. Behav. 53, 589–604. (doi:10.1006/anbe.1996.0334)

16. Zuberbühler K. 2000 Referential labelling in Diana monkeys. Anim. Behav. 59, 917–927. (doi:10.1006/anbe.1999.1317)

17. Zuberbühler K, Jenny D, Bshary R. 1999 The predator deterrence function of primate alarm calls. Ethology 105, 477–490. (doi:10.1046/j.1439-0310.1999.00396.x)

18. Isbell LA, Bidner LR. 2016 Vervet monkey (Chlorocebus pygerythrus) alarm calls to leopards (*Panthera pardus*) function as a predator deterrent. Behaviour 153, 591–606. (doi:10.1163/1568539X-00003365)

19. Wich SA, Nunn C. 2002 Do male ‘long-distance calls’ function in mate defense? A comparative study of long-distance calls in primates. Behav. Ecol. Sociobiol. 52, 474–484. (doi:10.1007/s00265-002-0541-8)

20. Snowdon CT. 2004 Sexual selection and communication. Cambridge: Cambridge University Press. (doi:10.1002/evan.10023)

21. Zuberbühler K. 2002 Effects of Natural and Sexual Selection on the Evolution of Guenon Loud Calls. In The Guenons: Diversity and Adaptation in African Monkeys. Developments in Primatology: Progress and Prospects. (eds ME Glenn, M Cords), pp. 289–306. Springer, Boston, MA. (doi:https://doi.org/10.1007/0-306-48417-X_20)

22. van Schaik CP, Bshary R, Wagner G, Cunha F. 2022 Male anti-predation services in primates as costly signalling? A comparative analysis and review. Ethology 128, 1–14. (doi:10.1111/eth.13233)

23. Fischer J, Kitchen DM, Seyfarth RM, Cheney DL. 2004 Baboon loud calls advertise male quality: acoustic features and their relation to rank, age, and exhaustion. Behav. Ecol. Sociobiol. 56, 140–148. (doi:10.1007/s00265-003-0739-4)

24. Kitchen DM, Seyfarth RM, Fischer J, Cheney DL. 2003 Loud calls as indicators of dominance in male baboons (*Papio cynocephalus ursinus*). Behav. Ecol. Sociobiol. 53, 374–384.

25. Mitani JC, Nishida T. 1993 Contexts and social correlates of long-distance calling by male chimpanzees. Anim. Behav. 45, 735–746. (doi:10.1163/1568539X-00003259)

26. Riede T, Arcadi AC, Owren MJ. 2007 Nonlinear acoustics in the pant hoots of common chimpanzees (*Pan troglodytes*): Vocalizing at the edge. J. Acoust. Soc. Am. 121, 1758–1767. (doi:10.1121/1.2427115)

27. Neumann C, Assahad G, Hammerschmidt K, Perwitasari-Farajallah D, Engelhardt A. 2010 Loud calls in male crested macaques, *Macaca nigra*: a signal of dominance in a tolerant species. Anim. Behav. 79, 187–193. (doi:10.1016/j.anbehav.2009.10.026)

28. McGregor PK, Peake TM. 2000 Communication networks: social environments for receiving and signalling behaviour. Acta Ethol. 2, 71–81.

29. Vehrencamp SL. 2000 Animal Signals: Signalling and Signal Design in Animal Communication. In Animal Signals: Signalling and Signal Design in Animal Communication (eds Y Espmark, T Amundsen, G Rosenqvist), pp. 277–300. Trondheim: Tapir Academic Press.

30. Benítez ME, le Roux A, Fischer J, Beehner JC, Bergman TJ. 2016 Acoustic and Temporal Variation in Gelada (Theropithecus gelada) Loud Calls Advertise Male Quality. Int. J. Primatol. 37, 568–585. (doi:10.1007/s10764-016-9922-0)

31. Fuller JL, Cords M. 2017 Multiple functions and signal concordance of the pyow loud call of blue monkeys. Behav. Ecol. Sociobiol. 71. (doi:10.1007/s00265-016-2230-z)

32. Fuller JL, Cords M. 2020 Versatility in a loud call: Dual affiliative and agonistic functions in the blue monkey boom. Ethology 126, 10–23. (doi:10.1111/eth.12955)

33. Zuberbühler K. 2006 Alarm calls. In Encyclopedia of Language & Linguistics (Second Edition) (ed K Brown), pp. 143–155. Oxford: Elsevier. (doi:10.1007/978-3-540-68706-1_3033)

34. Cheney DL, Seyfarth RM. 1981 Selective Forces Affecting the Predator Alarm Calls of Vervet Monkeys. Behaviour 76, 25–60.

35. Seyfarth RM, Cheney DL. 1985 Vervet Monkey Alarm Calls: Manipulation Through Shared Information? Behaviour 94, 150–166. (doi:10.1163/156853985X00316)

36. Henzi SP, Lucas J. 1980 Observations on the inter-troop movement of adult vervet monkeys (*Cercopithecus aethiops*). Folia Primatol. 33, 220–235. (doi:10.1159/000155936)

37. Young C, McFarland R, Ganswindt A, Young MMI, Barrett L, Henzi SP. 2019 Male residency and dispersal triggers in a seasonal breeder with influential females. Anim. Behav. 154, 29–37. (doi:10.1016/j.anbehav.2019.06.010)

38. Minkner MMI, Young C, Amici F, McFarland R, Barrett L, Grobler JP, Henzi SP, Widdig A. 2018 Assessment of Male Reproductive Skew via Highly Polymorphic STR Markers in Wild Vervet Monkeys, *Chlorocebus pygerythrus*. J. Hered. (doi:10.1093/jhered/esy048)

39. Price T, Ndiaye O, Hammerschmidt K, Fischer J. 2014 Limited geographic variation in the acoustic structure of and responses to adult male alarm barks of African green monkeys. Behav. Ecol. Sociobiol. 68, 815–825. (doi:10.1007/s00265-014-1694-y)

40. Price T, Wadewitz P, Cheney DL, Seyfarth RM, Hammerschmidt K, Fischer J. 2015 Vervets revisited: A quantitative analysis of alarm call structure and context specificity. Sci. Rep. 5, 13220. (doi:10.1038/srep13220)

41. Cohen J. 1960 A Coefficient of Agreement for Nominal Scales. Educ. Psychol. Meas. 20, 37–46. (doi:10.1177/001316446002000104)

42. Neumann C, Duboscq J, Dubuc C, Ginting A, Irwan AM, Agil M, Widdig A, Engelhardt A. 2011 Assessing dominance hierarchies: Validation and advantages of progressive evaluation with Elo-rating. Anim. Behav. 82, 911–921. (doi:10.1016/j.anbehav.2011.07.016)

43. R Core Team. 2021 R: A language and environment for statistical computing. *R Found. Stat. Comput. Vienna*, Austria. (doi:10.1016/j.jhevol.2005.09.003)

44. Bates D, Mächler M, Bolker B, Walker S. 2015 Fitting Linear Mixed-Effects Models using lme4. J. Stat. Softw. 67, 1–48. (doi:10.18637/jss.v067.i01)

45. Brooks ME, Kristensen K, van Benthem KJ, Magnusson A, Berg CW, Nielsen A, Skaug HJ, Mächler M, Bolker BM. 2017 glmmTMB balances speed and flexibility among packages for zero-inflated generalized linear mixed modeling. R J. 9, 378–400. (doi:10.32614/rj-2017-066)

46. Barr DJ, Levy R, Scheepers C, Tily HJ. 2013 Random effects structure for confirmatory hypothesis testing: Keep it maximal. J. Mem. Lang. 68, 255–278. (doi:10.1016/j.jml.2012.11.001)

47. Matuschek H, Kliegl R, Vasishth S, Baayen H, Bates D. 2017 Balancing Type I error and power in linear mixed models. J. Mem. Lang. 94, 305–315. (doi:10.1016/j.jml.2017.01.001)

48. Baayen RH. 2008 Analyzing linguistic data: A practical introduction to statistics using R. (doi:10.1017/s0305000909990080)

49. Schielzeth H. 2010 Simple means to improve the interpretability of regression coefficients. Methods Ecol. Evol. 1, 103–113. (doi:10.1111/j.2041-210x.2010.00012.x)

50. Fox J, Weisberg S. 2019 An {R} Companion to Applied Regression. Third. Thousand Oaks, CA: Sage publications.

51. Gelman A, Hill J. 2007 Data analysis using regression and multilevel/hierarchical models. Cambridge, UK: Cambridge university press.

52. Forstmeier W, Schielzeth H. 2011 Cryptic multiple hypotheses testing in linear models: overestimated effect sizes and the winner’s curse. Behav. Ecol. Sociobiol. 65, 47–55. (doi:10.1007/s00265-010-1038-5)

53. Dobson AJ, Barnett AG. 2018 An introduction to generalized linear models. Chapman and Hall/CRC.

54. Nieuwenhuis R, te Grotenhuis M, Pelzer B. 2012 Influence.ME: Tools for detecting influential data in mixed effects models. R J. 4, 38–47. (doi:10.32614/rj-2012-011)

55. McCullagh P, Nelder JA. 1989 Generalized Linear Models, 2nd Edn. (doi:10.2307/2347392)

56. Andelman SJ. 1987 Evolution of concealed ovulation in vervet monkeys (*Cercopithecus aethiops*). Am. Nat. 129, 785–799. (doi:10.1086/284675)

57. Harris TR. 2006 Within-and among-male variation in roaring by black and white colobus monkeys (*Colobus guereza*): What does it reveal about function? Behaviour 143, 197–218. (doi:10.1163/156853906775900702)

58. Searcy WA, Nowicki S. 2005 The evolution of animal communication: Reliability and deception in signaling systems. Princeton University Press.

59. Henzi SP, Bonnell T, Pasternak GM, Freeman NJ, Dostie MJ, Kienzle S, Vilette C, Barrett L. 2021 Keep calm and carry on: reactive indifference to predator encounters by a gregarious prey species. Anim. Behav. 181, 1–11. (doi:10.1016/j.anbehav.2021.08.024)

60. Teichroeb JA, Sicotte P. 2010 The Function of male agonistic displays in ursine colobus monkeys (*Colobus vellerosus*): Male competition, female mate choice or sexual coercion? Ethology 116, 366–380. (doi:10.1111/j.1439-0310.2010.01752.x)

61. Enquist M. 1985 Communication during aggressive interactions with particular reference to variation in choice of behaviour. Anim. Behav. 33, 1152–1161. (doi:10.1016/S0003-3472(85)80175-5)

62. Molles LE, Vehrencamp SL. 2001 Songbird cheaters pay a retaliation cost: Evidence for auditory conventional signals. Proc. R. Soc. B Biol. Sci. 268, 2013–2019. (doi:10.1098/rspb.2001.1757)

63. Fischer J, Hammerschmidt K, Cheney DL, Seyfarth RM. 2002 Acoustic features of male baboon loud calls: Influences of context, age, and individuality. J. Acoust. Soc. Am. 111, 1465–1474. (doi:10.1121/1.1433807)

64. Harris TR, Fitch WT, Goldstein LM, Fashing PJ. 2006 Black and white colobus monkey (*Colobus guereza*) roars as a source of both honest and exaggerated information about body mass. Ethology 112, 911–920. (doi:10.1111/j.1439-0310.2006.01247.x)

65. Erb WM, Hodges JK, Hammerschmidt K. 2013 Individual, contextual, and age-related acoustic variation in simakobu (*Simias concolor*) loud calls. PLoS One 8. (doi:10.1371/journal.pone.0083131)

66. Wheeler BC, Fischer J. 2012 Functionally referential signals: A promising paradigm whose time has passed. *Evol. Anthropol. Issues, News*, Rev. 21, 195–205. (doi:10.1002/evan.21319)

67. Ducheminsky N, Henzi SP, Barrett L. 2014 Responses of vervet monkeys in large troops to terrestrial and aerial predator alarm calls. Behav. Ecol. 25, 1474–1484. (doi:10.1093/beheco/aru151)

68. Price T, Fischer J. 2014 Meaning attribution in the West African green monkey: Influence of call type and context. Anim. Cogn. 17, 277–286. (doi:10.1007/s10071-013-0660-9)

69. Deshpande A, van de Waal E, Zuberbühler K. 2023 Context-dependent alarm responses in wild vervet monkeys. Anim. Cogn. (doi:10.1007/s10071-023-01767-0)

70. Behaviour A. 2018 Guidelines for the treatment of animals in behavioural research and teaching. Anim. Behav. 135, I–X. (doi:10.1016/j.anbehav.2017.10.001)

