## Supplementary material for "Alarm barks of male vervet monkeys as displays of male quality": Electronic supplementary material

### Supplementary figures and tables

Barking events were recorded throughout the home ranges of all six groups (fig. S1). When a barking event from one group was audible to a neighbouring group, outside of between group encounters, we occasionally estimated the transmission range of barks, by comparing GPS positions of the respective groups (fig. S2). While we only made few opportunistic recordings and could not account for weather, vegetation or caller elevation, we estimated barks to be audible over 900 m in some cases (fig. S2). The context of most events could not be determined (fig. S3). The distribution of the number of adult males that produced barks in calling events revealed that in the majority of events, only a single male called (fig. S4). Event durations showed considerable variation, with most events not lasting longer than about five minutes, while in rare cases durations of up to one hour were recorded (fig. S5).

The distribution of the number of barking events per month indicated an increase in monthly event frequencies during the mating season (fig. S6). The distribution of births during the study period suggested that most conceptions occurred in May in June, and only one apparently outside the mating season (fig. S7). Barking activity was highest around the time that most females conceived (fig. S6 and S7).

Since visual inspection of the first model (fig. 3) suggested that the effect of rank on barking probability appeared to depend strongly on the behaviour of the highest-ranked males, we fitted the model again but excluded the highest ranked males (fig. S8). The model without data from the highest-ranked males corroborated the assumption that the effect of rank on barking probability was strongly driven by the highest-ranked individuals (fig. S8). Plotting calling probability against male rank across all events, including those with unknown callers, suggested no sampling bias (fig. S9, for comparison see fig. 3 and S8).

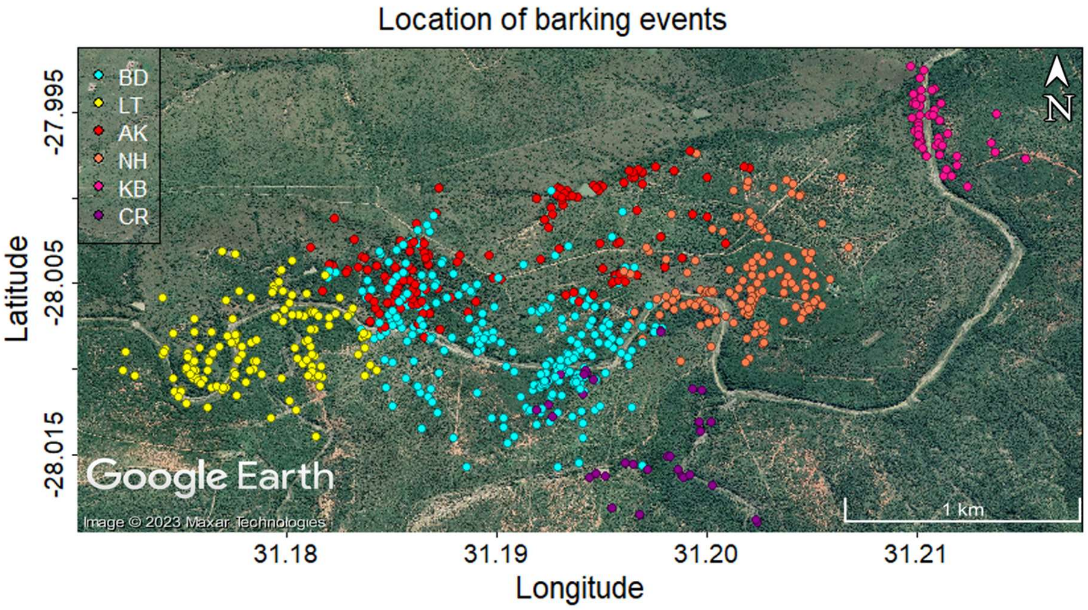


**Figure S1:** Location of barking events recorded from the six groups with available GPS data (N=840).

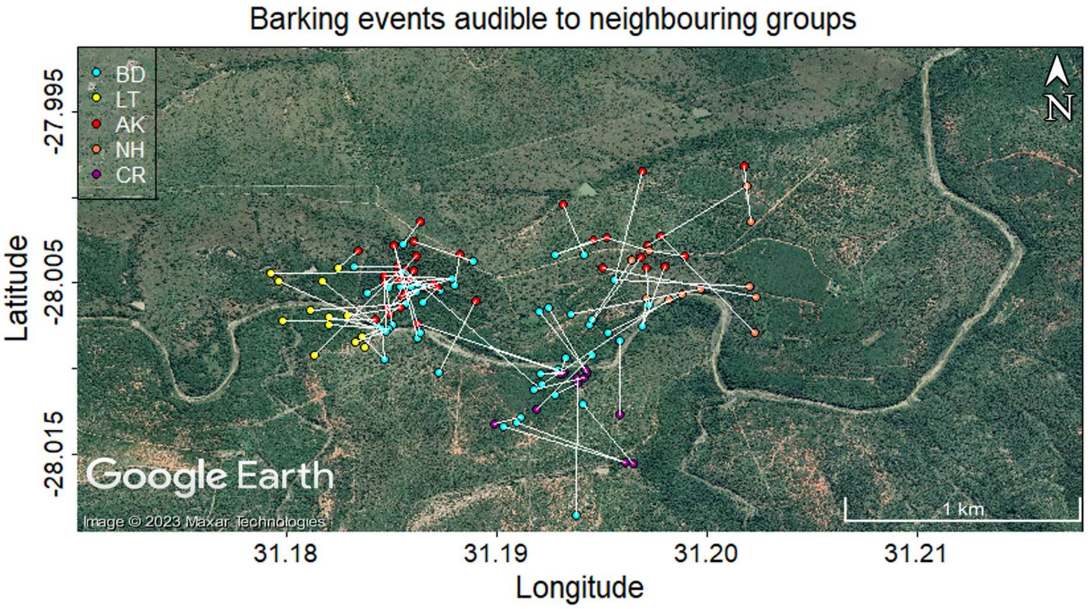


**Figure S2:** Barking event location relative to the location of a neighbouring group, at the time of the event, indicated by white lines. Recordings were made opportunistically when observers heard barks from another group outside of encounters (distance between groups > 100 m) and could confirm the barking group's identity via radio contact with observers present in the barking group (N=94).

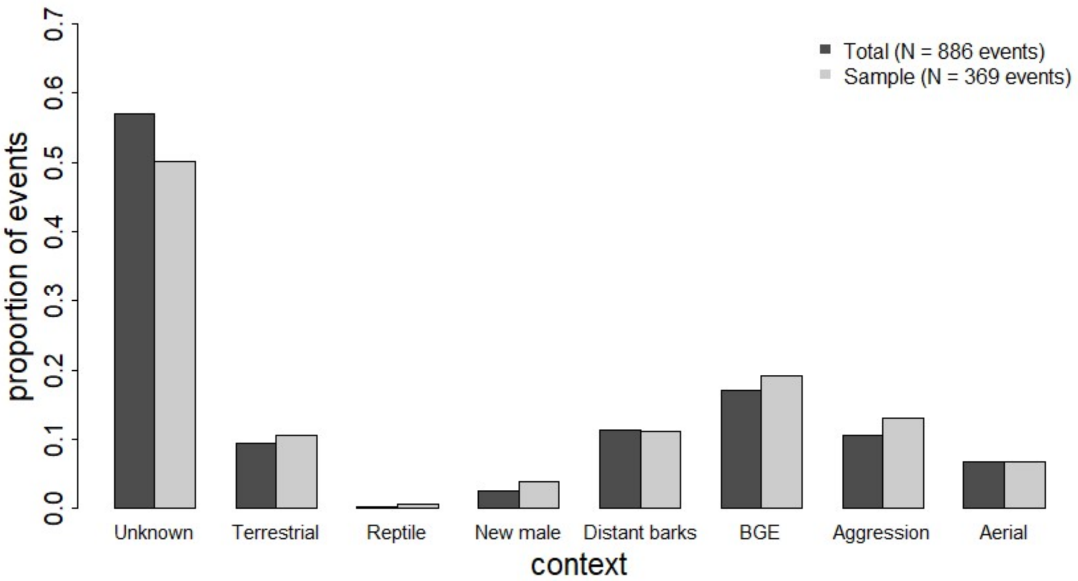


**Figure S3:** Relative proportions of contexts across all barking events (N=886, dark grey) and in the sample (N=369, light grey). Since barking events could be assigned to multiple contexts simultaneously, the plot shows the proportion of events a given context was scored in (context categories in table S5).

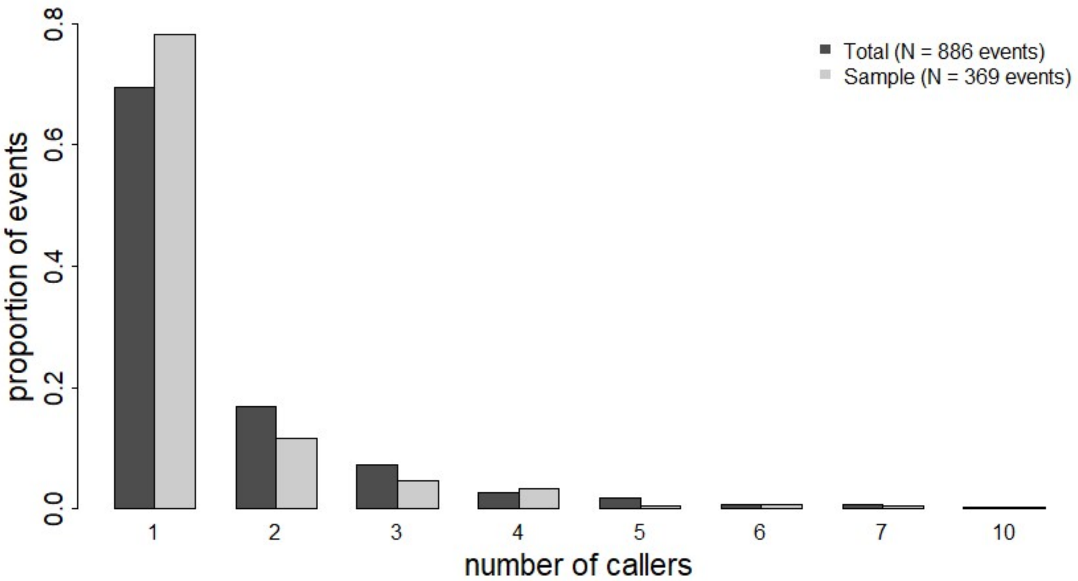


**Figure S4:** Distribution of the number of callers across all events (N=886, dark grey) and in the sample (N=369, light grey). In the majority of events, only a single adult male produced barks.

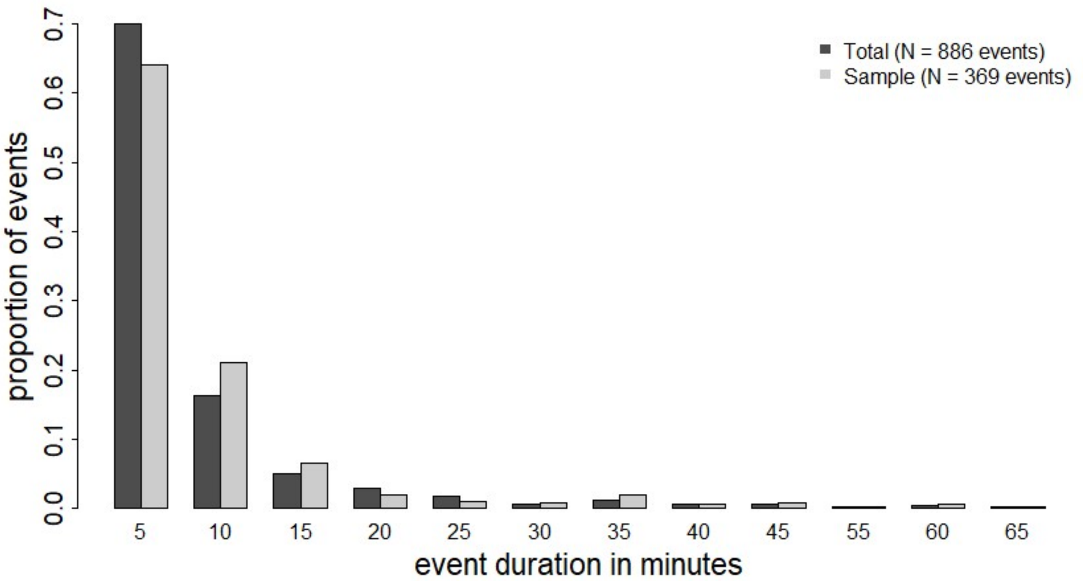


**Figure S5:** Distribution of event durations in 5 min intervals for all events (N=886, dark grey) and the sample (N=369, light grey). Most events last about 5 minutes.

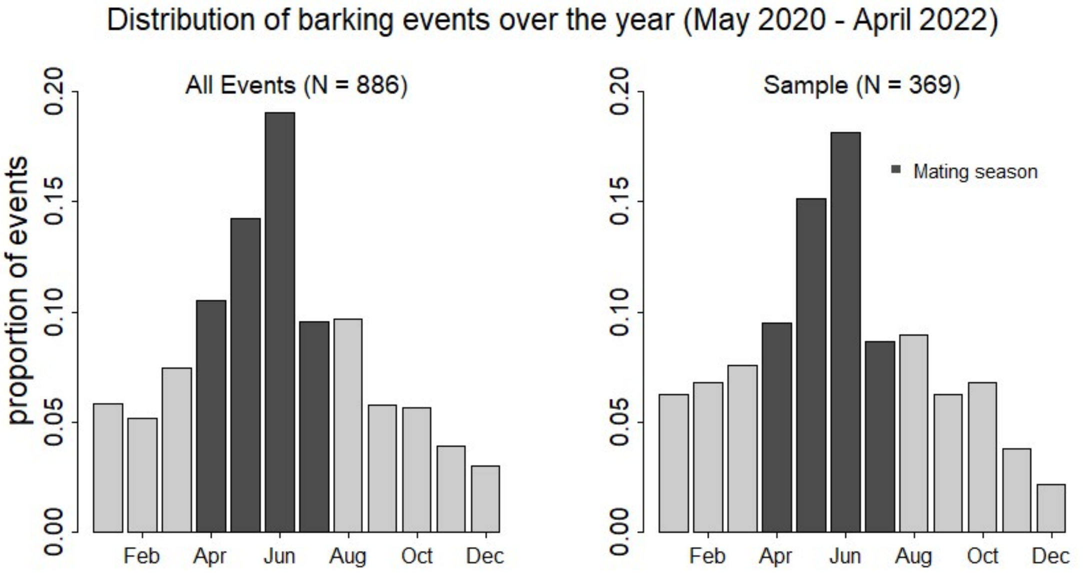


**Figure S6:** Proportion of calling events per month with mating season indicated.

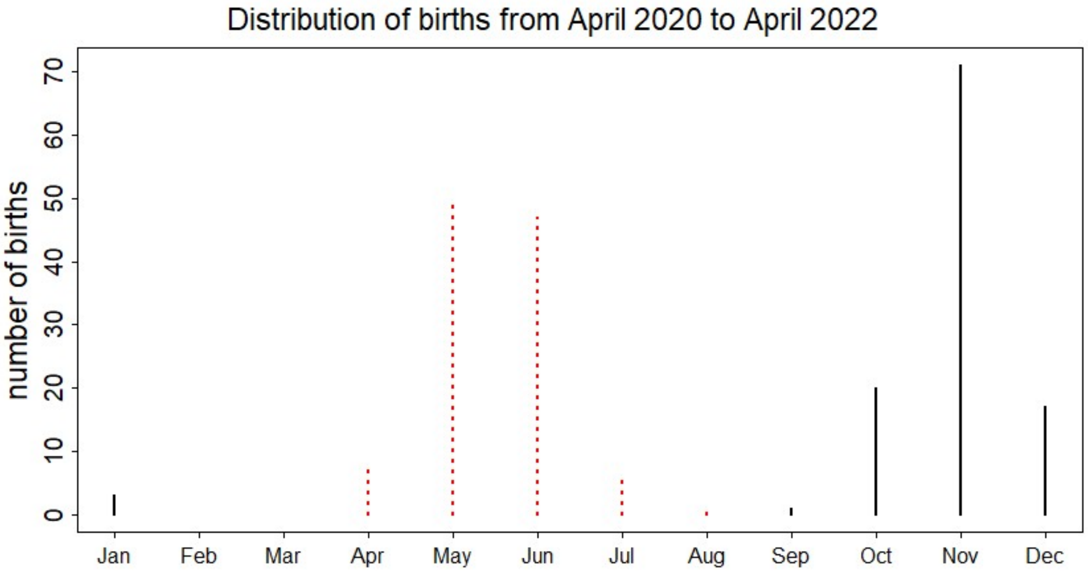


**Figure S7:** Total monthly number of births (black lines) and dated back times of conception (dashed red lines) in all six groups for the two year study period (N=112 births). The majority of births occurred in November. With a gestation period of 165 days, most conceptions occurred in May and June.

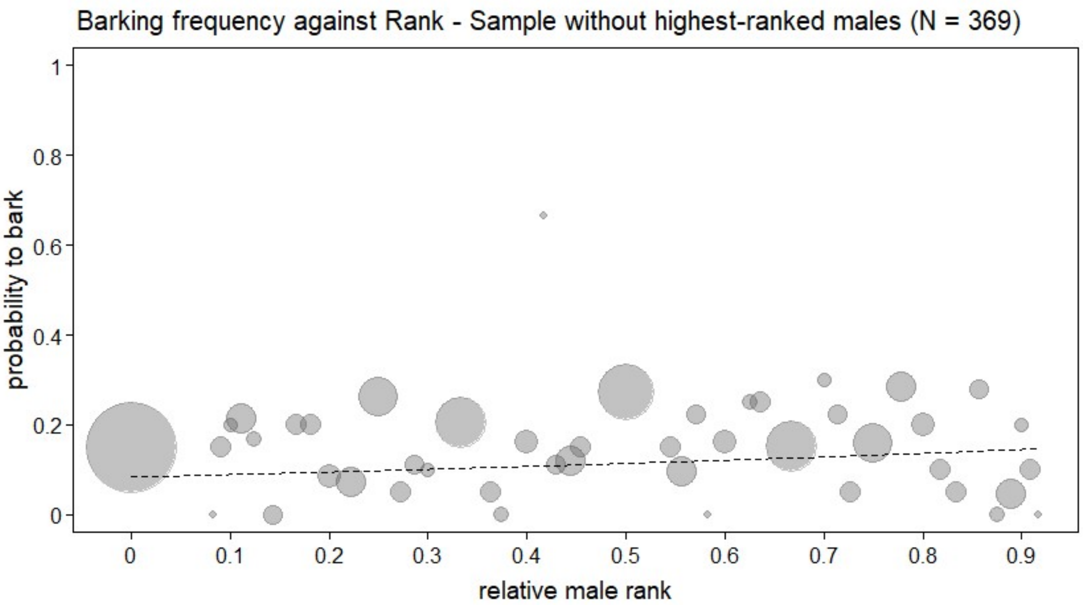


**Figure S8:** The effect of individual rank on barking probability excluding the highest ranked males (Rank = 1; 23 of 45 males temporarily held the highest rank). The effect of rank was likely driven by barking activity of the highest ranked males (Estimate = 0.202, Std. error = 0.166, z value = 1.215)

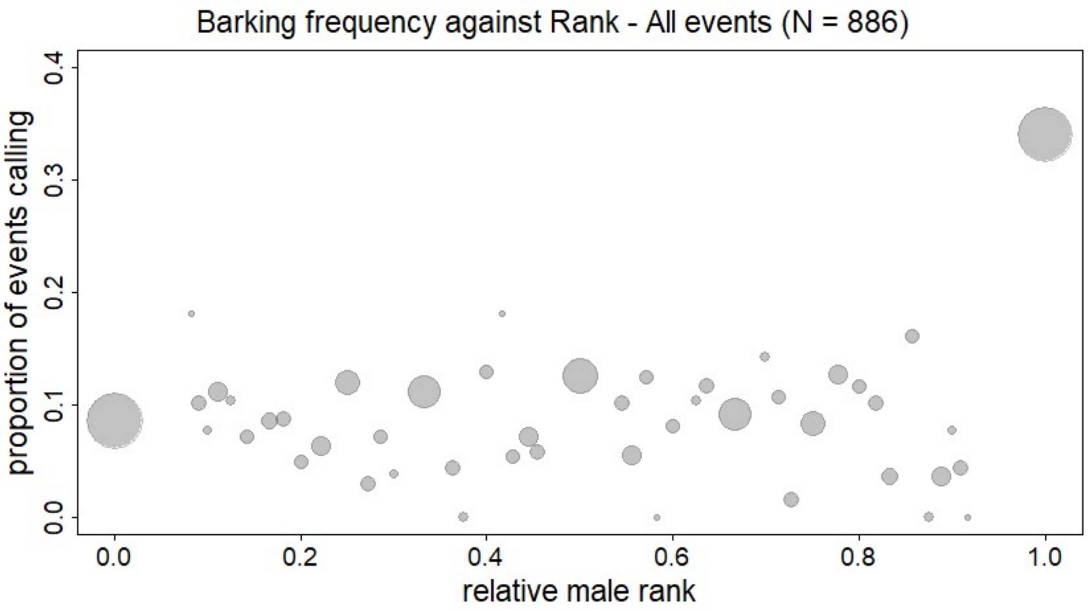


**Figure S9:** Barking probability against male rank across all events, including events with unknown callers.

### Supplementary tables

### Table S1: Model 1 – Individual calling probability in barking events – Full model

Model formula:

```
glmer(Bark ~ z.Rank*z.Males*MatingSeason + z.Rank*z.SexRatio*MatingSeason +z.Tenure+
    (1+ z.Rank*z.Males*MatingSeason.1 + z.Rank*z.SexRatio*MatingSeason.1 +z.Tenure ||Individual)+
    (1+ z.Rank*z.Males*MatingSeason.1 + z.Rank*z.SexRatio*MatingSeason.1 +z.Tenure ||Group)+
            (1 + z.Rank |Event_ID)+
            (1 + z.Rank ||Date.in.group),
data = t.data, family=binomial, control=glmerControl(optimizer="bobyqa", optCtrl = list(maxfun=1000000)))
```

Results of binomial model on barking probability. Shown are model estimates, standard errors, confidence intervals (CI), the test results obtained from likelihood ratio tests ($\chi 2$, *df*, *P*) and the range of estimates obtained when dropping levels of grouping factors one at a time (Min, Max). All covariates (Rank, Males, SexRatio and TenureMS) were z-transformed to a mean of 0 and a standard deviation of 1. The factor MatingSeason was dummy coded and centred in the random effects part, with the reference category set to outside MatingSeason.

| Term | Estimate | Std. Error | 2.5% CI | 97.5% CI | $\chi 2$ | *df* | *P* | Min | Max |
|---|---|---|---|---|---|---|---|---|---|
| Intercept | -1.818 | 0.261 | -2.28 | -1.325 | (-) | (-) | (-) | -1.934 | -1.5 |
| z.Rank | 0.75 | 0.234 | 0.306 | 1.166 | (-) | (-) | (-) | 0.453 | 1.61 |
| z.Males | -1.063 | 0.268 | -1.576 | -0.504 | (-) | (-) | (-) | -1.285 | -0.945 |
| MatingSeasonY | -0.047 | 0.207 | -0.463 | 0.367 | (-) | (-) | (-) | -1.082 | 0.037 |
| z.SexRatio | 0.028 | 0.17 | -0.312 | 0.343 | (-) | (-) | (-) | -0.07 | 0.109 |
| z.Tenure | 0.267 | 0.188 | -0.108 | 0.639 | 1.91 | 1 | 0.167 | 0.101 | 0.711 |
| z.Rank * z.Males | 0.064 | 0.236 | -0.4 | 0.508 | (-) | (-) | (-) | -0.094 | 0.979 |
| z.Rank * MatingSeasonY | -0.168 | 0.329 | -0.711 | 0.466 | (-) | (-) | (-) | -0.5 | 0.024 |
| z.Males * MatingSeasonY | 0.286 | 0.227 | -0.192 | 0.768 | (-) | (-) | (-) | -0.743 | 0.424 |
| z.Rank * z.SexRatio | 0.1 | 0.145 | -0.161 | 0.396 | (-) | (-) | (-) | -0.084 | 0.295 |
| MatingSeasonY * z.SexRatio | -0.098 | 0.237 | -0.546 | 0.403 | (-) | (-) | (-) | -0.332 | 0.042 |
| z.Rank * MatingSeasonY * z.Males | -0.073 | 0.332 | -0.657 | 0.515 | 0.048 | 1 | 0.827 | -0.475 | 0.101 |
| z.Rank * MatingSeasonY * z.SexRatio | -0.11 | 0.222 | -0.552 | 0.29 | 0.25 | 1 | 0.616 | -0.267 | 0.05 |

Model sample: N = 2055, Distribution of response: Bark (Yes) = 524, Bark (No) = 1531

Grouping factors: Individual (N=45), Group (N=6), Event ID (N=369), Date in Group (N=291)

**Table S2: Model 1 – Individual calling probability in barking events – Results of likelihood ratio tests assessing the two-way interactions**

| Term | *χ2* | *df* | *P* |
|---|---|---|---|
| z.TenureMS | 1.871 | 1 | 0.171 |
| z.Rank * z.Males | 0.025 | 1 | 0.874 |
| z.Rank * MatingSeason | 0.21 | 1 | 0.647 |
| z.Males * MatingSeason | 1.101 | 1 | 0.294 |
| z.Rank * z. SexRatio | 0.244 | 1 | 0.622 |
| MatingSeason * z. SexRatio | 0.223 | 1 | 0.637 |

**Table S3: Model 2 – Bark events per day – Full model**

Model formula:

glmmTMB(BarkEvents ~ MatingSeason * (z.Males + z.SexRatio) + z.GroupSize +
offset(log.TimeSpent) +
(1 + MatingSeason.1 * (z.Males + z. SexRatio) + z. GroupSize ||Group),
data = t.data, family=poisson, ziformula = ~1)

Results of the Poisson model on the number of barking events per day. Shown are model estimates, standard errors, confidence intervals (CI), the test results obtained from likelihood ratio tests ($\chi2$, *df*, *P*) and the range of estimates obtained when dropping levels of grouping factors one at a time (Min, Max). All covariates (Males, SexRatio, GroupSize) were z-transformed to a mean of 0 and a standard deviation of 1. The factor MatingSeason was dummy coded and centred in the random effects part, with the reference category set to outside MatingSeason.

| Term | Estimate | Std. Error | 2.5% CI | 97.5% CI | $\chi2$ | *df* | *P* | Min | Max |
|---|---|---|---|---|---|---|---|---|---|
| Intercept | -2.647 | 0.089 | -2.843 | -2.489 | (-) | (-) | (-) | -2.814 | -2.572 |
| MatingSeasonY | 0.712 | 0.090 | 0.531 | 0.886 | (-) | (-) | (-) | 0.65 | 0.884 |
| z.Males | 0.255 | 0.145 | -0.002 | 0.503 | (-) | (-) | (-) | -0.121 | 0.309 |
| z.SexRatio | 0.169 | 0.138 | -0.032 | 0.369 | (-) | (-) | (-) | 0.053 | 0.268 |
| z. GroupSize | -0.126 | 0.11 | -0.341 | 0.083 | 1.342 | 1 | 0.247 | -0.185 | -0.042 |
| MatingSeasonY * z.Males | -0.091 | 0.097 | -0.283 | 0.096 | 0.957 | 1 | 0.328 | -0.161 | 0.173 |
| MatingSeasonY * z.SexRatio | -0.243 | 0.146 | -0.493 | 0 | 2.4 | 1 | 0.121 | -0.36 | -0.105 |
| Zi@Intercept | -0.55 | 0.145 | -0.881 | -0.325 | (-) | (-) | (-) | -0.644 | -0.480 |

Model sample: N = 1915, Grouping factors: Group (N=6), Dispersion parameter: 1.08

**Table S4:** Definitions for age categories.

| Age category | Definition |
|---|---|
| **Adult female** | Females were considered adults as soon as they were four years old, which is the typical age of first reproduction. |
| **Adult male** | Males were considered adults as soon as they dispersed for the first time, which typically happens when they are four to five years old. |

**Table S5:** Descriptions of context categories that adult males responded to by producing barks. Researchers could assign multiple contexts for the same event if applicable.

| Context | Description |
|---|---|
| **Aerial** | Any aerial object, including bird species (predominantly raptors), occasionally helicopters, and small airplanes. |
| **Aggression** | Within group conflicts. |
| **Between group encounter (BGE)** | Encounters of one or more vervet monkey groups, defined as ranging within 100 meter of each another. |
| **Distant barks** | Barks from other groups faintly audible in the distance and outside of encounters. Note that in this dataset such events were only considered, if one or more males in the observed group responded to distant barks by producing barks themselves. Distant barking events without vocal responses from the observed group were not considered. |
| **Reptile** | Encounters with reptile species including mostly pythons, black mambas, spitting cobras, puff adders, spotted bush snakes and monitor lizards. |
| **Terrestrial** | Potential terrestrial threat, including mammalian land predators but also running antelopes and warthogs. Confirmed predators present at the site include leopards and caracals (seen on camera traps). Potential but unlikely predators include jackals and poaching dogs (observed while following monkeys). Note that predators are notoriously hard to confirm, meaning that if researchers could not identify any clear stimulus responsible for the calls, such events had to be classified as unknown (see below). |
| **New male** | Sighting of an unknown adult male in the group. |
| **Unknown** | Any barking events whose context could not be clearly determined. Note that this may include all the above categories and potential displays of males. Although male behaviour occasionally appeared to suggest a display, it could never be excluded that human observers had missed a predator or aggressive context. We therefore did not include ‘male display’ as a category since it could never have been scored without high uncertainty. Potential displays may involve shaking branches, jumping into a tree and barking from an elevated position while other monkeys seemed unconcerned. However, such behaviour also occurred in confirmed predator cases and aggressive interactions. |
